# Optimizing bioreactor growth of the smallest eukaryote

**DOI:** 10.1101/291211

**Authors:** Chuck R. Smallwood, Eric A. Hill, William Chrisler, Jory Brookreson, James E. Evans

## Abstract

Photosynthetic organisms are adept at circumventing nutrient deprivation. Microalgae in particular present novel adaptations to nutrient and light starvation since they can scavenge external and internal nutrient pools to redistribute energy resources for survival. In this report, a turbidostatic photobioreactor was used to characterize environmental conditions and nutrient requirements for cultures of the smallest free-living eukaryote *Ostreococcus tauri*. Optimized growth conditions were identified that enable 4-times faster phototrophic growth-rates while increasing total biomass 10-fold. By achieving phototrophic doubling times shorter than 6 hours, these results highlight the potential of this smallest eukaryote for future industrial bioproduct applications.

## Introduction

*Ostreococcus tauri* was originally isolated from copper contaminated water and it can grow in a variety of media conditions including artificial salt water (ASW) brackish media^1,2^ or with additives such as glycerol^3^. It has a highly condensed genome^4,5^, a simple cell ultrastructure^6^, is genetically tractable^7,8^, and is capable of significant triacylglycerol accumulation under various nutrient limitation conditions^9^. Additionally, it displays other favorable characteristics that can facilitate scale-up in bioreactors or downstream bioprocessing such as immotility, absence of a cell wall, and lack of annotated flocculating enzymes. Early reports of *O. tauri* isolates from natural marine environments described fast specific growth rates of 4.23 divisions per day^2^ and cell densities up to 2×10^10^ cell/mL^2,10^. Unfortunately, subsequent studies have not reported similar levels of sustainable fast *O. tauri* cell growth. Instead, while 2×10^8^ cells/mL can be achieved, a recent paper shows growth plots that take ~ 2 weeks to achieve similar yields with doubling times of ~4 days at high cell densities under analogous media, temperature and light conditions with original isolates^11^. Consequently, the slow native growth-rate of *O. tauri* in standard media and standard culturing methods leads to low biomass levels that would hinder cost effective industrial applications. Here, nutrient bioavailability, temperature, pH and light conditions for *O. tauri* were screened inside a custom photobioreactor to evaluate whether its growth could be enhanced to better meet the demands for downstream industrial applications.

## Results

Typical *O. tauri* growth conditions in the literature consist of standard or similar K media, incubation at room temperature (20–22°C) and a 12 to 16-hour alternating light:dark cycle with 20 *µ*E with white or blue light^12,13^. However, *O. tauri* can tolerate variable light intensities, spectral color profiles and different light:dark cycle ratios^14^. Using the average conditions with a conventional algae chamber and ventilated flasks, *O. tauri* cultures typically maxed out ~ 0.12 at OD730 (3×10^7^ cells/mL) with a doubling time of ~1.5 days. Because blue light is a high energy light source that could damage cellular components and effectively slow growth, illumination with lower energy per photon red light (680nm) was tested for cellular growth response. Cultures were grown with 360º light in a custom turbidostat bioreactor^15^ equipped with a built-in sensor to track dissolved oxygen accumulation as a metric of photosynthetic activity. Photosynthetic activity versus total red light fluence shows a slowed response from *O. tauri* in the bioreactor above illumination at 165 *µ*E/m^2^/s (Supp. Fig. 2). To avoid potential adverse impacts to cell health from constant illumination at max fluence, and to extend the practical life-time of LED light sources, the illumination was capped at ~50% max (80 *µ*E/m^2^/s) for all subsequent experiments.

After confirming *O. tauri* growth under constant red-light illumination, the question arose as to whether its diurnal cycling was necessary or simply a response to inconsistent energy or nutrient availability. Using the turbidostat bioreactor with continuous red LED illumination at 680nm, *O. tauri*’s diurnal cycles dampened in intensity and after 200 hours were eliminated (Fig. 1). Dissolved oxygen accumulation during the lighted phase indicated photosynthesis remained active even following the elimination of the diurnal cycle. Optimized growth rates were determined in real-time by modulating the turbidity of cultures via dilution at fixed incident light intensity to maintain growth at a relatively constant OD at 730nm (Supp. Fig. 2). At the beginning of the experiment the *O. tauri* culture had an average doubling time of 28 hours. Immediately after the diurnal cycle ceased, the doubling time was 17 hours but eventually stabilized at 11 hours as the culture came to steady-state within the turbidostat.

**Figure 1:**
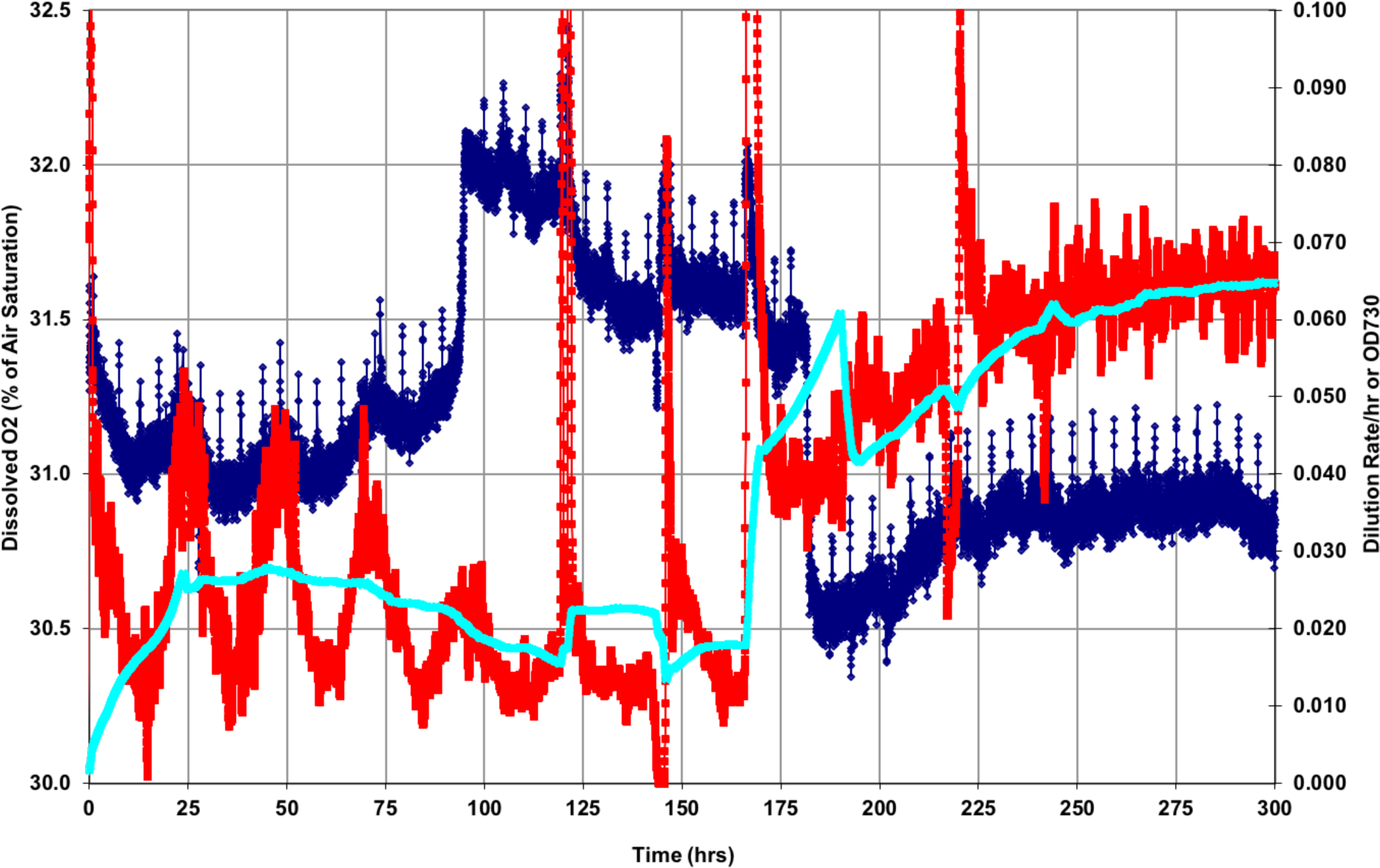
Photobioreactor elimination of diurnal rhythms and stabilization of growth. Simultaneous measurements of dissolved O2 (dark blue), dilution (red), and average dilution (cyan) were collected over 300 hours of *O. tauri* growth in the bioreactor under normal K media conditions and constant 680nm red light. Diurnal rhythms in *O. tauri* are evident between hour 0 through 175. Culture eventually stabilized beyond hour 200 in the bioreactor.

Despite the significant increase in growth rate following the removal of diurnal cycling, additional growth improvements were believed possible. This was supported by the observation that the growth phase in the turbidostat was exponential until an OD 730nm of 0.03 but declined above that threshold indicating light or nutrient limitation. Furthermore, the culture plateaued at an OD730 of 0.140, similar to culture flasks incubated in conventional growth chambers, suggesting that even with the improved doubling time, the total biomass was not significantly shifted. Subsequent optimization studies were therefore conducted below OD730 of 0.03 to maintain a stable baseline and compare the impact of additional supplementation of various media components on the overall growth rate.

Standard pH levels for K media are in the range 8.12 – 8.22^16^ and according to the NCMA website the recommended temperature ranges for *Ostreococcus* is 14 – 22ºC^17^ for laboratory cultures roughly mimic average oceanic conditions. The turbidostat was therefore used to evaluate the impact of varying temperature and pH on the culture growth in a reactor. *O. tauri* successfully grew at wider temperatures and pH ranges than previously reported. Limits for *O. tauri* cell viability were detected for temperatures above 34ºC and below 11ºC, and a maximal growth rate peaked ~ 28ºC (Fig. 2). For temperatures above 38ºC in the bioreactor, *O. tauri* is irreversibly damaged. However, cultures exposed to 35ºC for 12 hours eventually returned to full growth-rates after 53 hours. Using a similar approach and keeping the temperature constant at 28ºC, it was determined that growth rates struggle and fall off quickly below pH 7.5, whereas they stabilize above pH 7.5 (Fig. 3). For growth in a photobioreactor, a lower pH is convenient as it minimizes the likelihood of inorganic precipitation or clogging of sparge lines. Indeed, some precipitate was detected at pH levels above 8.2 in the bioreactor. Thus, while the cells can grow at the typical pH ~8.15, the optimal pH to enable maximal growth and minimize clogging due to inorganic precipitation in the turbidostat was pH 7.6.

**Figure 2:**
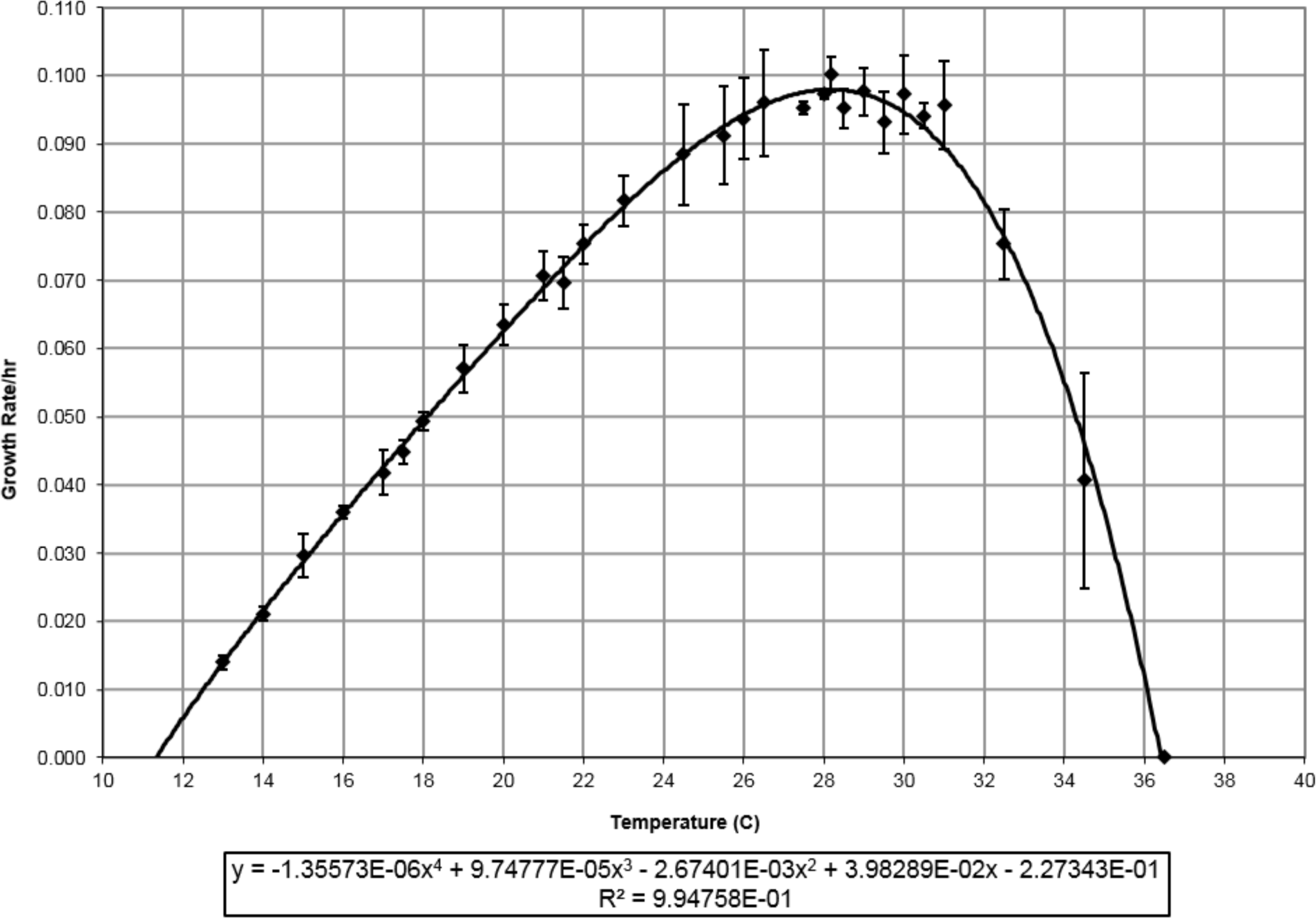
*O. tauri* can tolerate a wider range of temperatures than previously reported in the literature. Growth is observed at temperatures below 18°C, with an optimal growth rate at 28°C and negative exponential impacts above 30°C. A fourth order equation fits the data well, with an R-squared value of 0.993.

**Figure 3:**
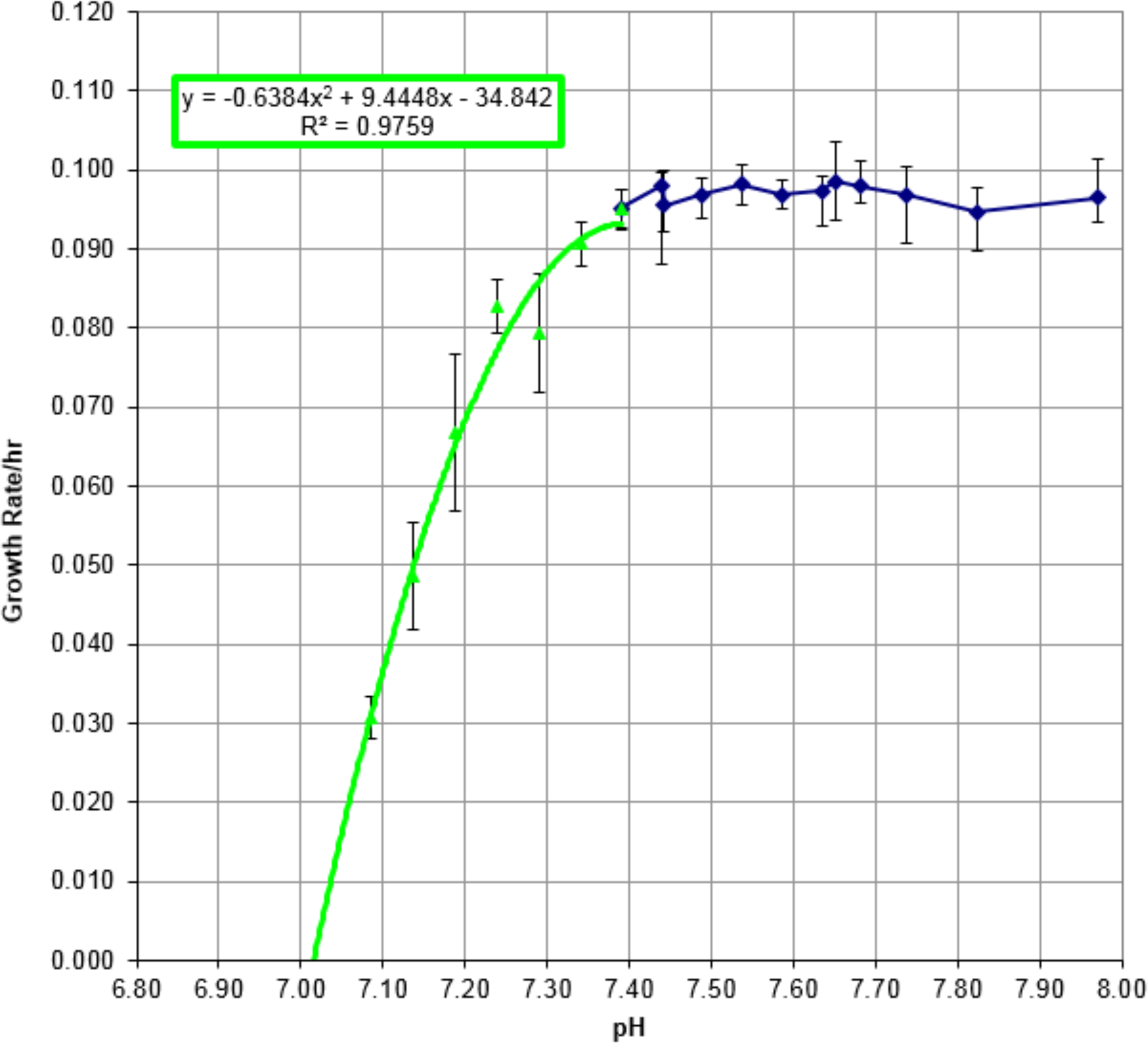
pH–dependent growth rate in the 680nm photobioreactor. Cellular growth rate per hour was measured at various culture pH at 28ºC in standard K media under 680nm red light in the photobioreactor. pH tolerance was repeated in both the decreasing (green) and increasing (blue) directions to ensure pH dependent effects on cellular growth rate. Growth rate remains optimal as long as the pH is above pH 7.5. Cells exposed to pH levels above 7.5 resulted in optimal growth rates, whereas declining growth rates are observed pH below 7.5.

After optimizing temperature (28°C) and pH (7.6) to optimized levels, the chemical composition of the media was adjusted to identify nutrient limitations. Standard K media combines ASW with both trace metals and vitamins (B12, thiamine and biotin). Based on the KEGG annotated genome of *O. tauri*, the cells should be able to synthesize and metabolize both thiamine and biotin, however vitamin B12 is clearly essential^18^. Surprisingly, injection of additional B12 into the turbidostat culture failed to cause a significant change in growth rate suggesting it was not the limiting factor. Similarly, injection of additional thiamine and biotin also failed to improve growth. However, supplementation of primary nutrients such as nitrogen and phosphorous did show significant changes.

Contrary to another report^10^, addition of ammonium chloride caused an immediate decrease in growth-rate (Supp. Fig. 3). This result could be due to ammonia gas formation or as others have reported in other algae^19^ it could be a result of intracellular toxicity from excess uptake by ammonium transporters. The other common nitrogen source in K media is nitrate. Other groups have reported the ability to maintain cultures of *Ostreococcus* solely containing nitrate as the nitrogen source at similar K media base concentrations (8.8×10^−4^M)^10,11,20^. Supplementation of additional sodium nitrate to 17 mM final concentration within the bioreactor not only caused elevated growth but also resulted in the rate remaining at maximum until about OD 0.080 at 730 nm (Supp. Fig. 4).

The photobioreactor was then reset to test the effects of phosphorus supplementation alone and then simultaneously with nitrogen supplementation. Utilizing 3-fold higher beta-glycerophosphate (bGP) with normal nitrogen levels, growth was elevated and the growth rate remained maximal until OD 0.171 @ 730 nm with the culture finally reaching a growth capacity of OD730 of 0.292 (Supp. Fig. 5). This was more than 2x the final attainalbe OD of the original culture in conventional growth chambers. However, supplementing the photobioreactor with 17 mM sodium nitrate and 0.37 mM bGP simultaneously achieved an OD of over 0.8 at OD730 (Supp. Fig. 6). Since *O. tauri* lacks any annotated pathway for bGP metabolism^18^, and it is more expensive than inorganic counterparts, experiments were performed to test whether other sources of phosphate could be utilized. Supplementation with the same concentration of PO_4_ but using NaH_2_PO_4_ yielded higher biomass results (Fig. 4). With these conditions, increased growth was retained to a much greater OD730 of 1.49, at which point the growth rate became limited by the amount of light that could be delivered to the cells. With the peak conditions combined, a maximal growth rate at OD730 of 0.122/hour was achieved with concurrent increased cell biomass. This equated to the doubling time of ~5.682 hours for cell division at 28.175ºC, pH 7.6. Dry weight samples taken from the culture at an OD of 1.2 at 730nm revealed an ash-free dry weight of 0.61939 g/L +/- 0.00315.

**Figure 4:**
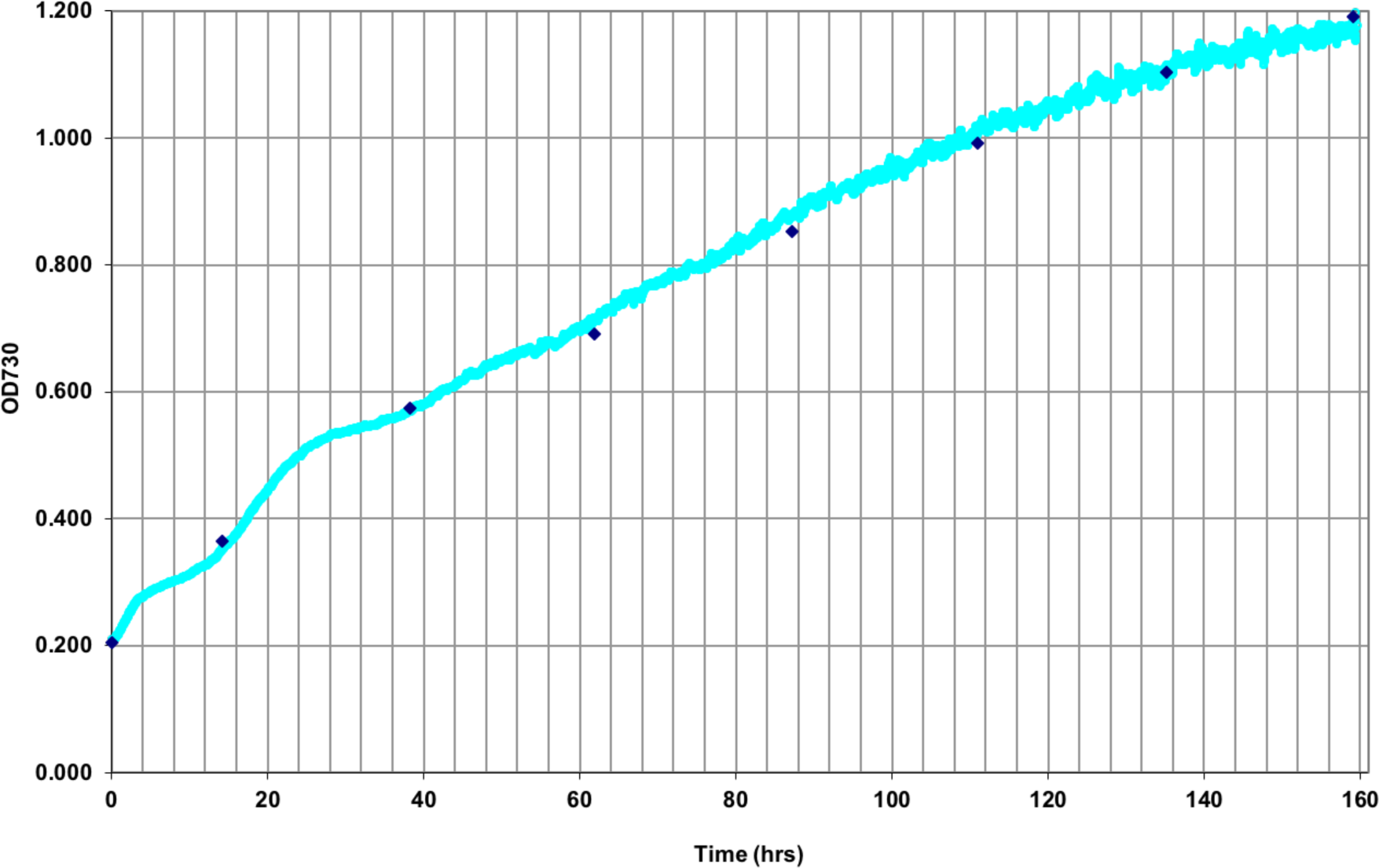
*O. tauri* growth in the photobioreactor under KOM conditions and red-light fluence. *O. tauri* growth was conducted with KOM conditions (K media with additions of 17 mM sodium nitrate and 0.37 mM NaH2PO4) at pH 7.6 in the photobioreactor with growth measured at OD730 estimate (cyan) with manually measured OD730 (blue diamonds). This experiment was not focused on maximum growth-rate, but on the maximal OD that could be obtained in the photobioreactor, which resulted in an OD730 above 1.2.

Lastly, after achieving improved growth rate and total biomass in the bioreactor, FACS enumeration and confocal imaging were used to verify whether cell morphologies changed. FACS enumeration showed more than 10-fold increases in microalgae cell concentration for cells in optimized nutrient conditions with a maximal cell density of 3×10^8^ cells/mL. This elevated cell density, if maintained in steady state with the improved doubling time of 6 hours, could achieve a total dry weight per illuminated area of 7.11 g/m^2^/day (Supp. Fig. 8), which is approaching the projected target of 18 g/m2/day proposed as the threshold for sustainable bioproduction^21^, which state the need for an average 18 g/m^2^/day.

Larger cell size distributions in populations found by FACS were further characterized by fluorescence confocal microscopy. Cells grown in standard conditions of low light (20*µ*E) and normal K media were typically less than 1*µ*m in cell diameter with spherical morphologies. However, photobioreactor optimized cells were about 2 times larger at 1.5 – 2*µ*m in diameter with spherical morphologies. Nile red staining of these larger cells revealed nominal neutral lipid content by confocal fluorescence microscopy and FACS (Fig. 5).

**Figure 5:**
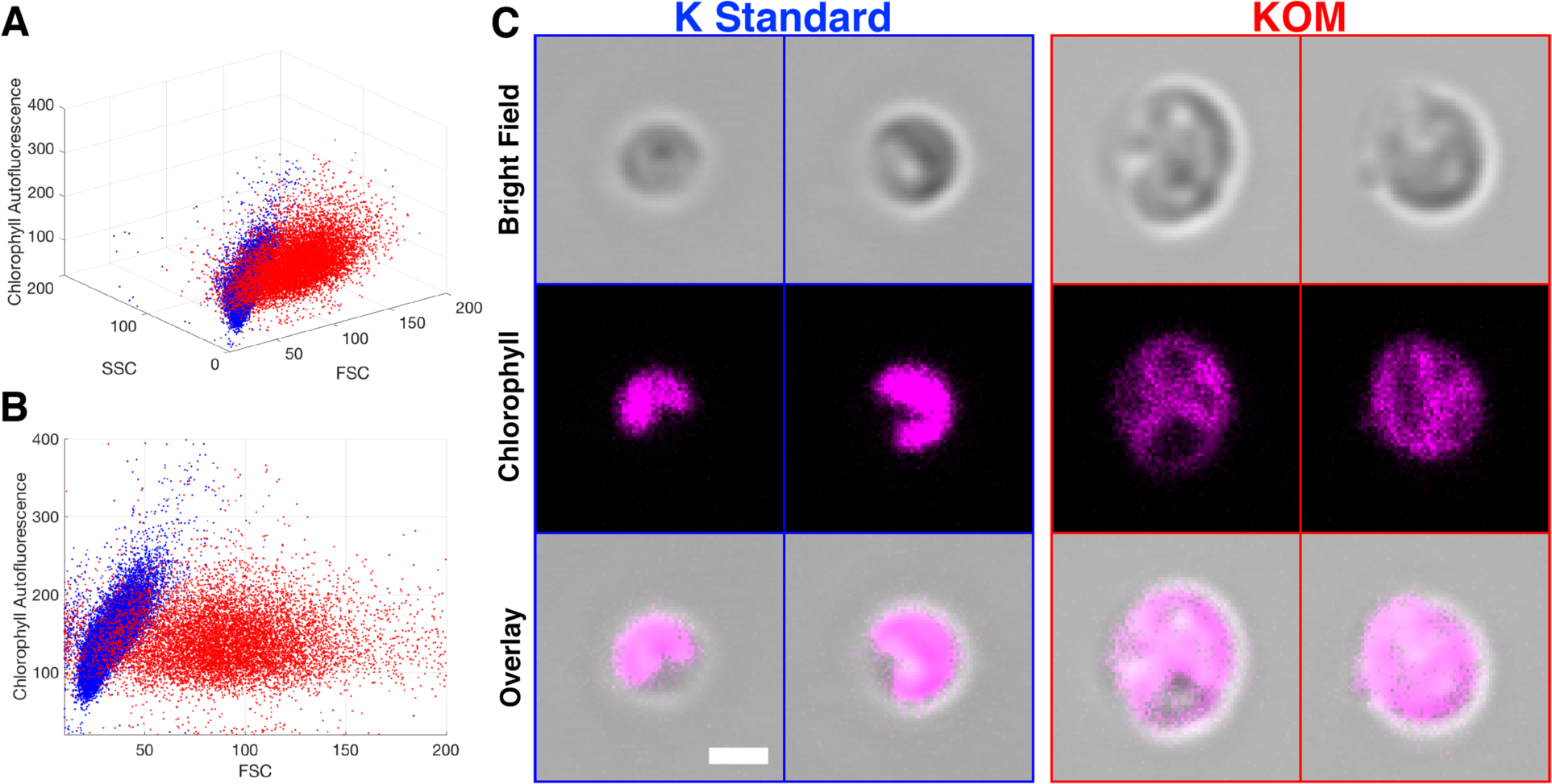
Single cell and population comparison of standard K media versus K optimized media (KOM) *O. tauri* cells grown under standard nutrient conditions with 20*µ*E blue light and K media (K Std) and cells grown in a red light photobioreactor with KOM conditions. Cells from standard K (blue) and KOM (red) were harvested and analyzed by FACS (A,B) for population dynamics. Cells were imaged by confocal fluorescence microscopy (C) to observe their chlorophyll (pink) autofluorescence, morphology, and cell size (C). Scale bar is 1*µ*m and equivalent for all images.

## Discussion

*O. tauri* are found in temperate to tropic to arctic oceans and throughout the photic zones^22–25^. The flexibility of this organisms to both thermal and light variations presents unique attributes that are ideal for a versatile, robust industrial bioprocessing strain. Here chemical and physical conditions were optimized for growing *O. tauri* in a red-light bioreactor to maximize photosynthetic efficiency and cell biomass for downstream industrial applications. Elimination of the diurnal cycle and growth under constant light were key to improving the growth-rate and allowed cultures to converge to steady-state growth. Supplementation with additional nitrate and phosphate improved cell doubling time and also increased total biomass capacity. Intriguingly, these results were attainable using wild-type *O. tauri* strains, which provides opportunities for further strain optimization by targeted genetic engineering that could possibly fine-tune *O. tauri* for even faster growth and elevated biomass. Overall, this study reveals that *O. tauri* is a small, resilient, and robust platform that warrants continued exploration for future bioprocessing applications. It also suggests that *O. tauri* subsists in nature on nutrient levels significantly lower than it is tuned for suggesting other native strains may also be environmentally limited.

## Methods

### Strains, photobioreactor settings, cell culture, and plating conditions

The strain OTTH0595 culture of *O. tauri* was obtained from the Roscoff Culture Collection (RCC745). Starter cultures were grown in standard Keller medium (K media) at 22°C with 25 *µ*E/m^2^/sec blue light from 17W T8 50/50 bulbs (Coralife, USA) fitted with 183 Moonlight Blue filters (Lee Filters, USA) on 12-hour light/dark cycles^1,13,16^. Batch-Mode Cultivation was conducted in a BioFlo 310 bioreactor (New Brunswick Scientific) modified with light sensors^15^ to maintain constant 680nm transmitted light intensities, 7.5 L vessel with a 5.5 L working volume, and a custom control loop to activate medium delivery pumps. The photosynthetic activity versus total 680nm red light fluence was conducted, which resulted in a plateau response above 165 *µ*E/m^2^/s illumination (Supp. Fig. 1). *O. tauri* cells were grown in the bioreactor under constant illumination with 680 nm LEDs. K media trace metals and vitamin mixes were added by filtration to autoclaved ASW and then covered with aluminum foil to prevent photo-oxidation of vitamin B12. Growth pH was initially 8.17 to mimic standard K media and average ocean pH. An air mixture of nitrogen and carbon dioxide was introduced by separate mass-flow controllers to achieve a final mix that was analyzed to be 0.42% CO_2_ by volume, 5.28% O_2_ by volume, with nitrogen at a total flow rate of 4L/min. For continuity, the gas mixture was left constant throughout all experiments. A 5.5L working volume of K media was used to cultivate *O. tauri* at temperatures from 13 to 36.5 °C, or from pH ~7 to 9.5. Harvesting of cells was conducted by pumping culture through sterile tubing with a peristaltic pump into 400 mL aliquots before centrifugation at 3,000 x g for 20 minutes at 22C. Sodium hydroxide (2M) and HCl (2M) were used to control pH throughout the experiment. Hydrochloric acid was added by a custom glass port through the head plate to avoid contact with the stainless-steel head plate.

For growth on plates, 2X artificial seawater (ASW) was made up with nutrients, vitamins, and trace metals added for either K standard media or Keller optimized media (KOM) conditions and then filter sterilized. Equal volumes of autoclaved molten 0.2% UltraPure^TM^ agarose (Thermo Fisher Scientific Inc., USA) was combined with sterile 2X solutions of K or KOM to produce 0.1% agarose semi-solid plates, respectively. *O. tauri* cells were harvested from a single mid-log culture grown under a 12-hour diurnal cycle in a blue light incubator in K standard media culture and centrifuged at 1000xG for 5 minutes to concentration cells to 10x before plating. Equal concentrations of cells were deposited onto the surface of K and KOM plates by pipet and incubated in a 680nm red light incubator with approximately 80*µ*E of constant illumination.

For dry weight determination, 400mL aliquots of cells were pelleted and suspended in purified water in pre-baked and pre-weighed aluminum weigh-boats. The cell pellet suspension was then dried at 100C, and then re-weighed. Finally, the dry pellets were combusted at 550C in a furnace and then re-weighed to determine the ash-free dry weight.

### Fluorescence activated cell sorting (FACS) analysis of photobioreactor cultures

*O. tauri* cells were cultured to mid log phase in the photobioreactor, harvested and fixed with paraformaldehyde at specific light, temperature, pH, or OD730 points and stored in 4ºC before each experimental measurement on the BD INFLUX flow cytometer (BD Biosciences, San Jose, CA, USA). Forward (FSC) and side (SSC) scatter were used to gate out any non-specific cellular debris and chlorophyll autofluorescence. Each individual FACS experiment was calibrated to 3.6 mm Ultra Rainbow Fluorescent Particles (Spherotech, Lake Forest, IL, USA) and maintained specific gate setting for relative analysis of each experiment. Data was plotted using Matlab.

### Confocal fluorescence and SRS microscopy

Photobioreactor cultured *O. tauri* cells were imaged on a Leica SP8 Confocal Microscope with HyD Detector using a 100× 1.4na water immersion objective with 1×, 2x and 4× zoom to satisfy Nyquist frequency. Auto-fluorescence of chlorophyll (red) was excited at 638nm, and emissions were monitored at 675–685 nm and then displayed over bright-field channel. Cells were immobilized on glass slides with poly-L-lysine and imaged immediately with *z*-scan slicing of 0.34*µ*m to survey whole cells. All channels maintained identical gain and laser power settings to across the time course to provide relative levels of fluorescent intensity. No adjustments of contrast or gain were applied to fluorescence channels during post processing.

### Calculation of total biomass from bioreactor

The fastest growth rates observed during this study were estimated in the OD730 growth region between 0.01 – 0. Thus, we assume for biomass calculations that if the culture was maintained in this high growth rate region of 0.12 (OD730), which is 10x fewer cells than the highest cell density of 1.2 (OD730), then the estimate Ash Free Dry Weight (AFDW) of 0.619g/L calculated from the KOM bioreactor culture at elevated cell densities. From the calculated total illuminated area in the bioreactor of 0.14 m2 and a volume of 5.5L, the whole reactor at 0.12 (OD730) would be 5.5L multiplied by 0.0619g/L to equal 0.34g total AFDW. Therefore, the total AFDW per illuminated area would be 0.34g divided by 0.14m2 to equal 2.43g/m2. Since the fastest growth rate obtain was 0.122(OD730) per day, the total AFDW per illuminated area per day estimate would be 2.43g/m2 multiplied by (0.122/hour) multiplied by 24hours/day equals 7.11g/m2/day.

## Acknowledgements

This research was performed using the Environmental Molecular Sciences Laboratory (EMSL), a national scientific user facility sponsored by the Department of Energy’s office of Biological and Environmental Research and located at Pacific Northwest National Laboratory (PNNL). This work was supported by DOE-BER Mesoscale to Molecules Bioimaging Project FWP #66382.

## References

1 Keller, M. D., Selvin, R. C., Claus, W. & Guillard, R. R. L. Media for the Culture of Oceanic Ultraphytoplankton. J Phycol 23, 633–638 (1987).

2 Courties, C. et al. Phylogenetic analysis and genome size of Ostreococcus tauri (Chlorophyta, Prasinophyceae). J Phycol 34, 844–849, doi:DOI 10.1046/j.1529-8817.1998.340844.x (1998).

3 Chuck R Smallwood, W. C., Jian-Hua Chen, Emma Patello, Mathew Thomas, Rosanne Boudreau, Axel Ekman, Hongfei Wang, Gerry McDermott, James E Evans. *Ostreococcus tauri* is a high-lipid content green algae that extrudes clustered lipid droplets. BioRXiv 249052, doi:https://doi.org/10.1101/249052 (2018).

4 Derelle, E. et al. Genome analysis of the smallest free-living eukaryote Ostreococcus tauri unveils many unique features. Proc Natl Acad Sci U S A 103, 11647–11652, doi:10.1073/pnas.0604795103 (2006).

5 Blanc-Mathieu, R. et al. An improved genome of the model marine alga Ostreococcus tauri unfolds by assessing Illumina de novo assemblies. BMC Genomics 15, 1103, doi:10.1186/1471-2164-15-1103 (2014).

6 Henderson, G. P., Gan, L. & Jensen, G. J. 3-D ultrastructure of O. tauri: electron cryotomography of an entire eukaryotic cell. PloS one 2, e749, doi:10.1371/journal.pone.0000749 (2007).

7 Lozano, J. C. et al. Efficient gene targeting and removal of foreign DNA by homologous recombination in the picoeukaryote Ostreococcus. Plant J 78, 1073–1083, doi:10.1111/tpj.12530 (2014).

8 Lelandais, G. et al. Ostreococcus tauri is a new model green alga for studying iron metabolism in eukaryotic phytoplankton. BMC Genomics 17, 319, doi:10.1186/s12864-016-2666-6 (2016).

9 Degraeve-Guilbault, C. et al. Glycerolipid Characterization and Nutrient Deprivation-Associated Changes in the Green Picoalga Ostreococcus tauri. Plant Physiol 173, 2060–2080, doi:10.1104/pp.16.01467 (2017).

10 Fouilland, E. et al. Productivity and growth of a natural population of the smallest free-living eukaryote under nitrogen deficiency and sufficiency. Microb Ecol 48, 103–110, doi:10.1007/s00248-003-2035-2 (2004).

11 Lupette, J. et al. Marinobacter Dominates the Bacterial Community of the Ostreococcus tauri Phycosphere in Culture. Front Microbiol 7, 1414, doi:10.3389/fmicb.2016.01414 (2016).

12 Moulager, M., Corellou, F., Verge, V., Escande, M. L. & Bouget, F. Y. Integration of light signals by the retinoblastoma pathway in the control of S phase entry in the picophytoplanktonic cell Ostreococcus. PLoS Genet 6, e1000957, doi:10.1371/journal.pgen.1000957 (2010).

13 O’Neill, J. S. et al. Circadian rhythms persist without transcription in a eukaryote. Nature 469, 554–558, doi:10.1038/nature09654 (2011).

14 Thommen, Q. et al. Probing entrainment of Ostreococcus tauri circadian clock by green and blue light through a mathematical modeling approach. Front Genet 6, 65, doi:10.3389/fgene.2015.00065 (2015).

15 Melnicki, M. R. et al. Feedback-controlled LED photobioreactor for photophysiological studies of cyanobacteria. Bioresour Technol 134, 127–133, doi:10.1016/j.biortech.2013.01.079 (2013).

16 Berges, J. A., Franklin, D. J. & Harrison, P. J. Evolution of an artificial seawater medium: Improvements in enriched seawater, artificial water over the last two decades. J Phycol 37, 1138–1145 (2001).

17 Sciences, B. L. f. O. Bigelow National Center for Marine Algae and Microbiota.

18 Kanehisa, M., Sato, Y., Kawashima, M., Furumichi, M. & Tanabe, M. KEGG as a reference resource for gene and protein annotation. Nucleic Acids Res 44, D457–462, doi:10.1093/nar/gkv1070 (2016).

19 Bittsanszky, A., Pilinszky, K., Gyulai, G. & Komives, T. Overcoming ammonium toxicity. Plant Sci 231, 184–190, doi:10.1016/j.plantsci.2014.12.005 (2015).

20 Djouani-Tahri el, B., Sanchez, F., Lozano, J. C. & Bouget, F. Y. A phosphate-regulated promoter for fine-tuned and reversible overexpression in Ostreococcus: application to circadian clock functional analysis. PloS one 6, e28471, doi:10.1371/journal.pone.0028471 (2011).

21 Productivity Enhanced Algae and Tool-Kits. Report No. DE-FOA-0001628, (DOE Office of Energy Efficiency and Renewable Energy, https://eere-exchange.energy.gov/, 2017).

22 Treusch, A. H. et al. Phytoplankton distribution patterns in the northwestern Sargasso Sea revealed by small subunit rRNA genes from plastids. ISME J 6, 481–492, doi:10.1038/ismej.2011.117 (2012).

23 Cheah, W. et al. Response of phytoplankton photophysiology to varying environmental conditions in the Sub-Antarctic and Polar Frontal Zone. PloS one 8, e72165, doi:10.1371/journal.pone.0072165 (2013).

24 Guillou, L. et al. Diversity of picoplanktonic prasinophytes assessed by direct nuclear SSU rDNA sequencing of environmental samples and novel isolates retrieved from oceanic and coastal marine ecosystems. Protist 155, 193–214, doi:10.1078/143446104774199592 (2004).

25 Lopes Dos Santos, A. et al. Diversity and oceanic distribution of prasinophytes clade VII, the dominant group of green algae in oceanic waters. ISME J 11, 512–528, doi:10.1038/ismej.2016.120 (2017).

